# NANO DOPA MELANIN PIGMENT WITH COSMETIC POTENTIAL PRODUCED BY HALOTOLERANT MARINE *CORYNEBACTERIUM AMYCOLATUM*

**DOI:** 10.64898/2026.08.14.744987

**Authors:** Shereen M Murshidah, Noble K Kurian, P Aiswarya, Sreeja Narayanan

## Abstract

Bacterial melanin are macromolecules found in nature that provide a wide range of biological functions, including pigmentation, resistance to radiation, scavenging of free radicals, thermoregulation and protected from oxidative stress and harmful heavy metals. The melanin is crucial for pathogenesis and bacterial survival in a variety of circumstances, and they can also influence how bacteria interact with other organisms. Usually, bacteria produce the melanin is either black or brown colour. The produced melanin has excellent properties like antimicrobial, antioxidant, photoprotective and antibiofilm. This is a report on *Corynebacterium amycolatum* melanin-producing bacteria isolated from the marine sediment of Thiruvanmiyur beach in Tamil Nadu, India. *Corynebacterium amycolatum* was screened using tyrosine basal broth (TBB), and UV-visible spectroscopy, FTIR, and SEM were used to analyse the extracted melanin. The non-pathogenic nature of the *Cornynebacterium amycolatum* strain was verified through antibiotic sensitivity profiling. The cosmetic potential was evaluated using antioxidant and SPF assays. *Corynebacterium amycolatum* predominantly uses the DOPA pathway for melanin production, was confirmed using kojic acid inhibitor study. The *in vitro* studies on mouse fibroblast cell line (L929) and *in vivo* studies on zebra fish embryos shows non-cytotoxicity using this melanin, even in lower concentration confirms its potential to use in cosmetic formulation. This research aims to demonstrate that bacterial melanin is safe for the environment and has qualities that make it safer and more effective in cosmetics.

## INTRODUCTION

Melanin are its hydrophobic pigments with a negative charge and high molecular weight. (Nosanchuk & Casadevall.,2003) and are widely distributed, showing significant role in a wide range of organisms. Many animals, plants, and certain fungi and bacteria possess melanin pigment, which are heterogeneous pigment formed through the oxidative polymerization of indolic or phenolic compounds (Pavan et al.,2020). Based on their structural monomers, melanin can be classified into three primary types: allomelanin, pheomelanin, and eumelanin. The polymer of eumelanin is mostly made up of indole-type units that are produced when L-tyrosine or L-DOPA (L3,4-dihydroxyphenylalanine) are oxidation. The oxidative polymerization of the cysteinyl conjugate of DOPA via benzothiazine intermediates leads to in pheomelanin as the final product. Cells have seemed like black or brown colored by eumelanin and allomelanin, whereas cells are yellow or red colored by pheomelanin, which is found in the animal kingdom. Pyomelanin and neuromelanin are two other forms of melanin. (Singh et al.,2025).

Many bacteria virulence and pathogenicity toward their specific animal or plant hosts have been associated with their capacity to produce melanin. Numerous helminthic, bacterial, and fungal diseases are caused by the microorganisms that may synthesize melanin pigments. Melanin role in virulence has been most thoroughly investigated in association with fungal diseases. The problem with melanin is that existing biochemical and biophysical methods cannot give this complex polymer a chemical structure since they are amorphous, insoluble, and not suitable for structural investigations using liquids or crystallography (Nosanchuk & Casadevall.,2003)

Melanin has found commercial use in cosmetics as a UV protection. Melanin protects the body from the sun due to its photoprotective property. Because melanin cannot breakdown easily, it can absorb up to 90% of the heat produced by sunlight. When exposed to UVA rays, melanin interacts with DNA and acts as a photosensitizer, producing reactive oxygen species (ROS). More importantly, melanin promotes the production of histamine, which helps fair-skin people experience sun induced redness and swelling. Because of its large number of unpaired electrons,vmelanin pigment effectively scavenges the reactive oxygen species, including free radicals. As an antioxidant property found in bacterial melanin that prevents free radicals via a sequence of one-electron transfer events, melanin may be used in cosmetics to reduce tissue damage caused by toxins (El-Zawawy et al.,2024). Naturally bacterial melanin has the same property as like chemical sunscreen, but the main advantage melanin does not provide any carcinogenic effect. So bacterial melanin was good alternative instead of chemically prepared sunscreen. The melanin produced by microorganisms has been shown to be affordable and environmentally sustainable and it is an alternative solution to chemically synthesized pigments (El-Zawawy et al.,2024).

This research mainly focused on the characterization of melanin-produced by bacteria *Corynebacterium amycolatum,* and exploring the pigments potential in cosmetic applications.

## MATERIALS AND METHODS

### SAMPLE LOCATION

The marine sediment sample was collected from Thiruvanmiyur Beach, Tamil Nadu, India. (12°.97’, 80°.26’) and the sample was carefully transferred into a sterile container. Then, this sample were serially diluted and spread plated to a nutrient agar plate that had 3% NaCl in it. The plates were incubated for 24 hours at 37°C. The colonies on the plates vary in color and the distinct morphology of the strains was used to get the pure culture. Quadrant streaking in a nutrient agar plate with 3% NaCl and a 24-hour incubation period was used to carry out for pure colony. The grown pure colonies were screened for the melanin production.

### SCREENING FOR MELANIN PRODUCING BACTERIA PRIMARY SCREENING

The isolated pure colonies were screened through TBB (tyrosine basal broth) (Antony et al.,2023). The TBB is the minimal media that contains KH2PO4, MgSO4, and NaCl. The isolated pure strains were inoculated into TBB, and the test tubes were kept in a shaking incubator for an incubation period of 15 days at 37°C. The colour will be changed in the media and monitored daily. After 15 days, the melanin-produced bacteria were scaled up into 100ml in tyrosine basal broth for secondary screening (Jigna et al.,2022).

### MELANIN PRODUCTION

The melanin produced bacteria was scaled up into 100ml of TBB (tyrosine basal broth) medium. Add 10% of inoculum to the TBB broth. The flask was kept in a shaking incubator for an incubation period of 15 days at 37°C. The colour will be changed in the media and monitored daily (Jigna et al.,2022).

### CHARACTERIZATION OF MELANIN PRODUCING BACTERIA CTB10 GRAM STAINING AND BIOCHEMICAL CHARACTERIZATION OF CTB10

Gram staining is used to determine the Gram nature of strain CTB10 (Choi.,2021). IMViC and catalase test were performed as per the protocol mentioned further. Catalase test is performed by taking a loop full of strain made a smear on the microscopic slide add one or two drops of hydrogen peroxide (H2O2), if catalase is present, oxygen bubbles will form. These are observations to determine the biochemical properties of the bacteria (Hemraj et al.,2013).

### MOLECULAR CHARACTERIZATION OF CTB10 STRAIN

The genomic DNA of the CTB10 strain was extracted, and its purity was assessed using agarose gel electrophoresis. The 16S rRNA were then amplified using PCR using the universal primers (forward primer: CAGGCCTAACACATGCAAGTC) and (reverse primer: GGGCGGWGTGTACAAGGC). The PCR product was then examined using agarose gel electrophoresis with 1.2% of agarose gel. ExoSAP-IT treatment is used to remove unwanted primers and dNTPs and the Bigdye terminator v3.1 cycle sequencing kit (Applied Biosystems, USA) was used in a PCR thermal cycler (Geneamp PCR system 9700, Applied Biosystems) according to manufacturer instructions. Finally, Geneious Pro v5.1 software was used to perform sequence analysis (Janda & Abbott., 2007).

### PHYLOGENETIC ANALYSIS OF CTB10 STRAIN

Phylogenetic tree was constructed using MEGA11 software (molecular evolutionary genetics analysis version 11) using the neighbour-joining method with bootstrapping of 1000 (Calvopiña et al.,2023).

### ANTIBIOTIC SENSITIVITY PROFILING

The antibiotic sensitivity profiling test of CTB10 was done Mueller Hinton agar plates by the disc diffusion method. Spread the broth culture uniformly on the MHA plate and place the antibiotic disc on the MHA plate and incubated for 24 hours at 37°C. Measure the zone formation around each antibiotic disc. The grams of each antibiotic disc are Streptomycin (S25)-25mcg, Gentamicin (GEN10)- 10mcg, Ampicillin (AMP2)-2mcg, Cefepime (CPM30)-30mcg, Penicillin(P10)-10units, cefoperazone (CFS75/30)-75/30mcg, Kanamycin(K30)-30µg and Ceftriaxone (CIT30/10)- 30/10mcg. (Hudzicki.,2009). The MAR index was determined using the formula a/b, where "a" stands for the number of antibiotics to which an isolate was resistant and "b" for the total number of antibiotics disc was tested (Mir et al.,2022).

### HALOTOLERANCE OF STRAIN CTB10

The halotolerance test is helpful to find whether bacteria may grow and survive in environments with high salt concentrations. Melanin-producing CTB10 strain was streaked on nutrient agar plates with different NaCl concentrations like 2%, 4%, 6%, 8%, 10%, and 15% NaCl and incubated for 24 hours at 37°C during the halotolerance test. The observation to determine the bacterial growth tolerance (Ilyas et al.,2020 & Rahman et al.,2017).

### EXTRACTION AND PURIFICATION OF MELANIN

The bacteria produced melanin in the tyrosine basal broth (TBB) media. Discard the pellet and keep the supernatant by using centrifugation at 5000 rpm and boiling the supernatant at 97°C, which allows the evaporation of excess water content and makes it condense. Then, perform acid precipitation by using 0.1N HCL and further washing procedures, including 0.1N HCL, 70% ethanol, and distilled water. These washing steps help to remove acid debris and some impurities, then the purified melanin is kept in the lyophilizer to remove the wet part from the bacterial melanin. The purified bacterial melanin was used for further studies (Ferraz et al., 2021).

### CHARACTERIZATION OF CTB10 MELANIN SOLUBILITY TEST

The CTB10 bacterial melanin is tested for its solubility, and it was tested by using the solvents, including water, ethanol, methanol, isopropanol, hydrochloric acid, sulfuric acid, DMSO, and sodium hydroxide. Take the melanin and vortex, centrifuge at 5000 rpm for 5 minutes. Observe the melanin, whether soluble or not (Jigna et al.,2022).

### UV- VISIBLE SPECTROPHOTOMETRY

The absoption of melanin in UV and visible wavelength were evaluated. 100µg/ml melanin solution in NaOH is used in the analysis, and the absorbance was measured between 200 to 700nm, and 0.1N NaOH was the blank solution of the analysis. (Pralea et al.,2019).

### FT-IR SPECTROPHOTOMETRY

FTIR analysis was used to profile the active functional groups present in the melanin (Rudrappa et al., 2022). The melanin sample combined with KBr, an FTIR spectrophotometer was used to perform the FTIR analysis (Tarangini & Mishra.,2013). The Infrared rays pass through the bacterial melanin and range from 4000-400cm-1, and the result was compared with earlier reports.

### FIELD EMISSION SCANNING ELECTRON MICROSCOPY (FE-SEM)

The morphological characteristics of the prepared melanin were examined using field emission scanning electron microscopy (Carl Zeiss Sigma FE-SEM, Germany) with a Schottky thermal field emitter as the electron source. The pigment was first centrifuged, dried, and mounted onto carbon tape. The samples were then transferred onto copper grids, coated with a thin layer of gold via sputtering, and imaged under the FE-SEM. Image acquisition and basic measurements were performed using the SMARTSEM software.

### COSMETIC PROPERTY OF CTB10 MELANIN

#### SUN PROTECTION FACTOR (SPF) ANALYSIS

Sun protection factor is helpful in finding the bacterial melanin SPF capability. SPF assay is done as per Mansur equation (Patki et al.,2021). 100µg/mL melanin in 1mL ethanol is used for the study. The blank was 100% ethanol for this analysis (Antony et al.,2023).

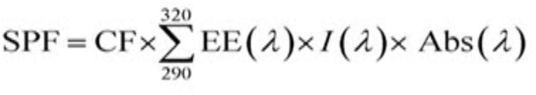

CF (correction factor) =10, EE (1) =arrhythmogenic impact of radiation with wavelength k, Abs (1) = spectrophotometric absorbance value of the solution, and I= solar intensity spectrum were the formulas used to determine SPFs. EE (1) ×I was found to be constant.

### ANTIOXIDANT ACTIVITY OF CTB10 MELANIN DPPH RADICAL SCAVENGING ASSAY

In DPPH assay the ascorbic acid was used as standard reference.1mg/ml ascorbic acid and same concentration of melanin was dilute with distilled water and 60µM of DPPH solution made freshly, then 200µl DPPH solution mixed with 50µg/ml melanin sample from this take 1.56, 3.12, 6.25, 12.5, 25, 50, 100, 200, 400,800 and 1000µl. This experiment was performed in a 96-well plate, and the plate was kept in the dark for 15 minutes at room temperature, and absorbance was measured at 515nm. The control is DPPH and a 95% methanol blank for this analysis (Brand-Williams et al.,1995 and Anthony & Saleh.,2013).

### *IN-VITRO* CYTOTOXICITY STUDY OF CTB10 MELANIN CELL LINE AND MAINTENANCE

The cell line of L929 was purchased from the National Centre for Cell Sciences (NCCS), Pune, India. The cells were cultured in Dulbecco’s modified eagle medium (DMEM-Himedia), supplemented with 10% heat inactivated fetal bovine serum (FBS) and 1% antibiotic cocktail containing penicillin (100 u/ml), streptomycin (0.1mg/ml) and amphotericin B (0.25 µg/ml) and the cell were culture in the TC flask and maintained at 37°C.

### MTT ASSAY

A 96-well plate was used for maintaining the cells, which were then cultured for 24 hours at 37°C with a 5% CO2. After mixing 10mg/ml of melanin with 10mg/ml of DMSO, the mixture was filtered and sterilized using a 0.2µm Millipore syringe filter. The DMEM media is used to dilute the melanin solution, and the melanin sample load on the 96-well plate with cell culture has concentrations of 6.25, 12.5, 25, 50 and 100µg/ml. The cells without melanin treatment are used as a control, without cells as a blank for this analysis, and the plates are incubated for 24 hours. This experiment was performed in triplicate to reduce result errors. After incubation, the media inside the well was discarded, and 100µl of MTT solution in PBS was added. Again, incubating for 2 hours allows the formation of formazan crystals. Centrifuge the content in the well plate and discard the supernatant. 100µl of DMSO is loaded into the well. Measure the absorbance at 570nm using a microplate reader (Mosmann.,1983, Joseph et al.,2012).

The cell viability formula:

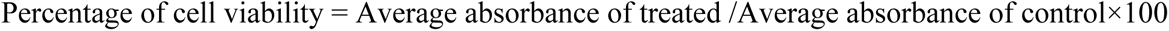

### *IN-VIVO* CYTOTOXICITY STUDY OF CTB10 MELANIN IN ZEBRAFISH (DANIO RERIO) EMBRYOS

The CTB10 melanin sample was tested with an animal model of an in-vivo study using Zebra fish (*Danio rerio*) embryos. The zebra fish embryos were bought from the aquarium and 1mg/mL CTB10 melanin was dissolved in NaOH, then the dissolved melanin was made neutral to pH 7. This experiment was carried out in 24-well plate. The diluted melanin sample was loaded into the 24 well plate in different concentrations (1µg, 3.125µg, 6.25µg, 12.5µg, 25µg, 50µg,75µg and 100µg). Each well five zebra fish embryos were added, and the embryos were grown in the tested CTB10 melanin solution dissolved in fish water. The embryos were observed in 24 hours and 48 hours through a microscope by checking their phenotypic variation and embryo viability. The embryos without melanin treatment were used as a control (Joshi & Katti.,2018).

Percentage of zebra fish embryo viability = No of live embryos in each concentration / total number of embryos in each concentration× 100

### STUDYING – MELANIN METABOLISM IN STRAIN CTB10 USING INHIBITOR

The inhibitor study was helpful in identifying the melanin biosynthesis pathway. The inhibitor study using kojic acid was done. Kojic acid 0.5 to 2mM were added to the melanin production media and incubated to observe the effect of inhibitor on melanin production. TBB without kojic acid was the control of the experiment. The test tubes were kept in a shaking incubator for an incubation period of 15 days at 37°C. Observe the test tube and measure the melanin concentration were found out spectrophotometrically (Manivasagan et al.,2013 & Choi et al., 2014).

## RESULT AND DISCUSSION

### ISOLATION OF MELANIN PRODUCING BACTERIA

Thirty-one colonies were isolated from the spread plate, and from these ten distinct colonies were selected based on colours, morphology of the bacteria, and streaked on quadrants to get pure colonies. Isolated pure colonies were used for screening for melanin production.

### SCREENING OF MELANIN PRODUCING BACTERIA PRIMARY SCREENING

The selected strains were screened in the TBB (tyrosine basal broth) media to check whether they produce melanin or not. The four strains produce the melanin in the TBB media, and the strains are CTB3, CTB5, CTB7 and CTB10 (Figure 1). The CTB10 strain produces the melanin on the 5^th^ day, producing melanin faster and good colour intensity. So, further research has been conducted using the CTB10 strain.

**Figure 1:**
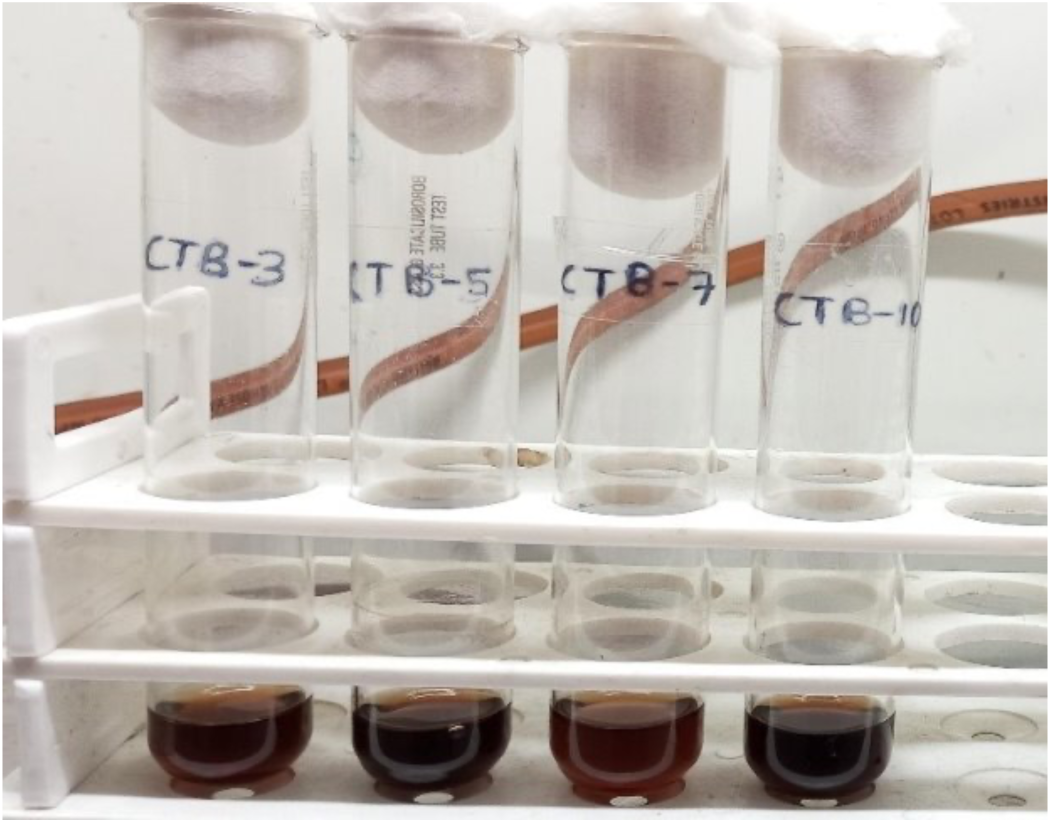
Primary screening-Melanin producing bacteria in tyrosine basal broth in the boiling tubes. The melanin producers are CTB3, CTB5, CTB7 and CTB10.

### SECONDARY SCREENING

The selected CTB10 strain was scaled up into 100ml of tyrosine basal broth media in a conical flask and the strain started melanin production on the 5^th^ day and completed melanin production on the 15^th^ day. The final colour of melanin was dark brown. The TBB media white colour during inoculation (Figure 2a) and 15^th^ day of melanin production (Figure 2b).

**Figure 2:**
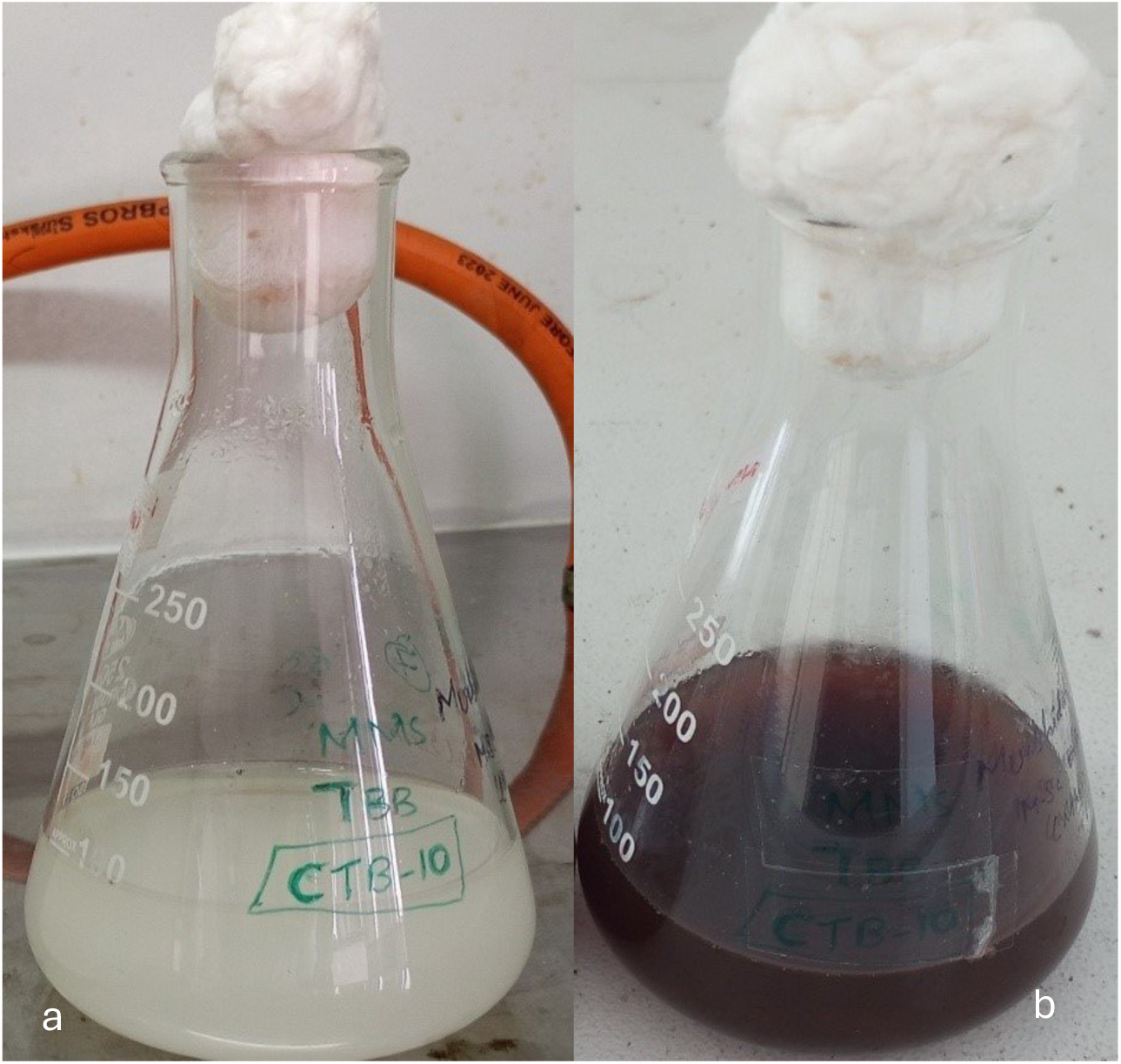
Secondary screening – melanin production by CTB 10 strain (a) Before inoculation; (b) Melanin production in 15^th^ day

### CHARACTERIZATION OF MELANIN PRODUCING BACTERIA CTB10 GRAM STAINING

The characterization of CTB 10 began with Gram staining, and it revealed the CTB10 strain Gram positive rod (Figure 3). The other melanin producing Gram positive rod bacteria are *Bacillus, Amorphotheca*, and *Streptomyces* (Choi.,2021).

**Figure 3:**
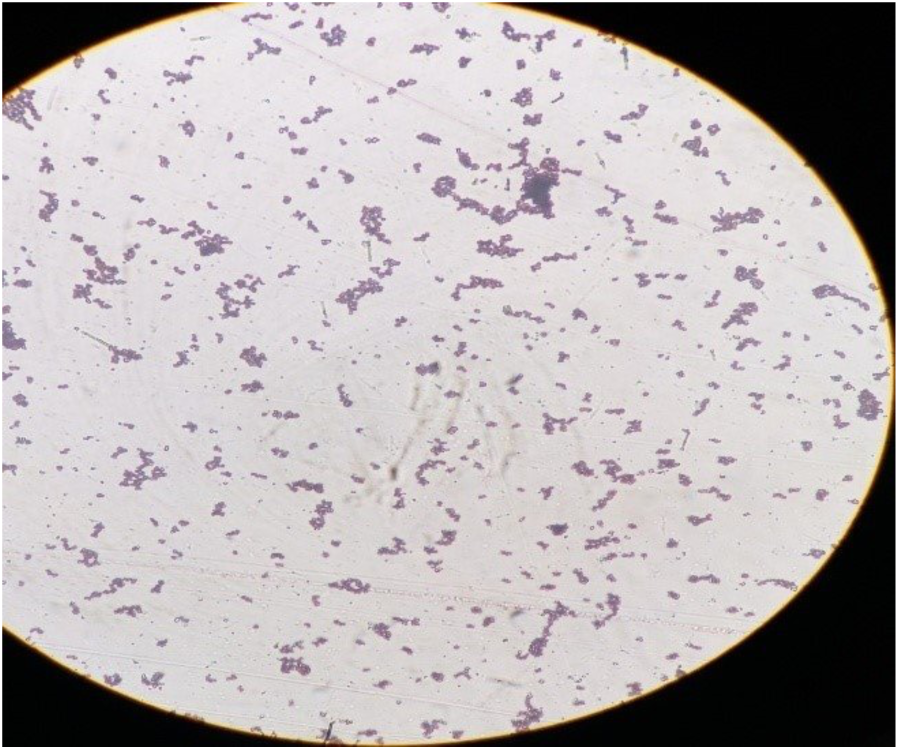
Gram staining of CTB10 strain – Gram positive rod

### BIOCHEMICAL CHARACTERIZATION OF CTB10

The biochemical characterization by IMViC and Catalase tests revealed CTB10 was found to be negative for Indole, Vogus Proskauer and Citrate and Methyl red was fond to be positive. CTB10 was found to produce catalase enzyme. (Table 1).

**Table 1:**
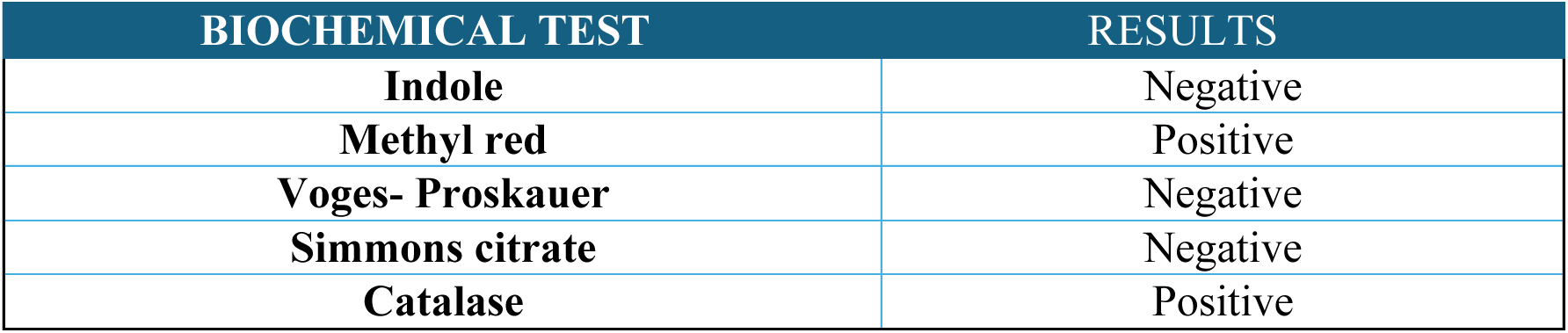
Biochemical tests for strain CTB10.

**Table 2:**
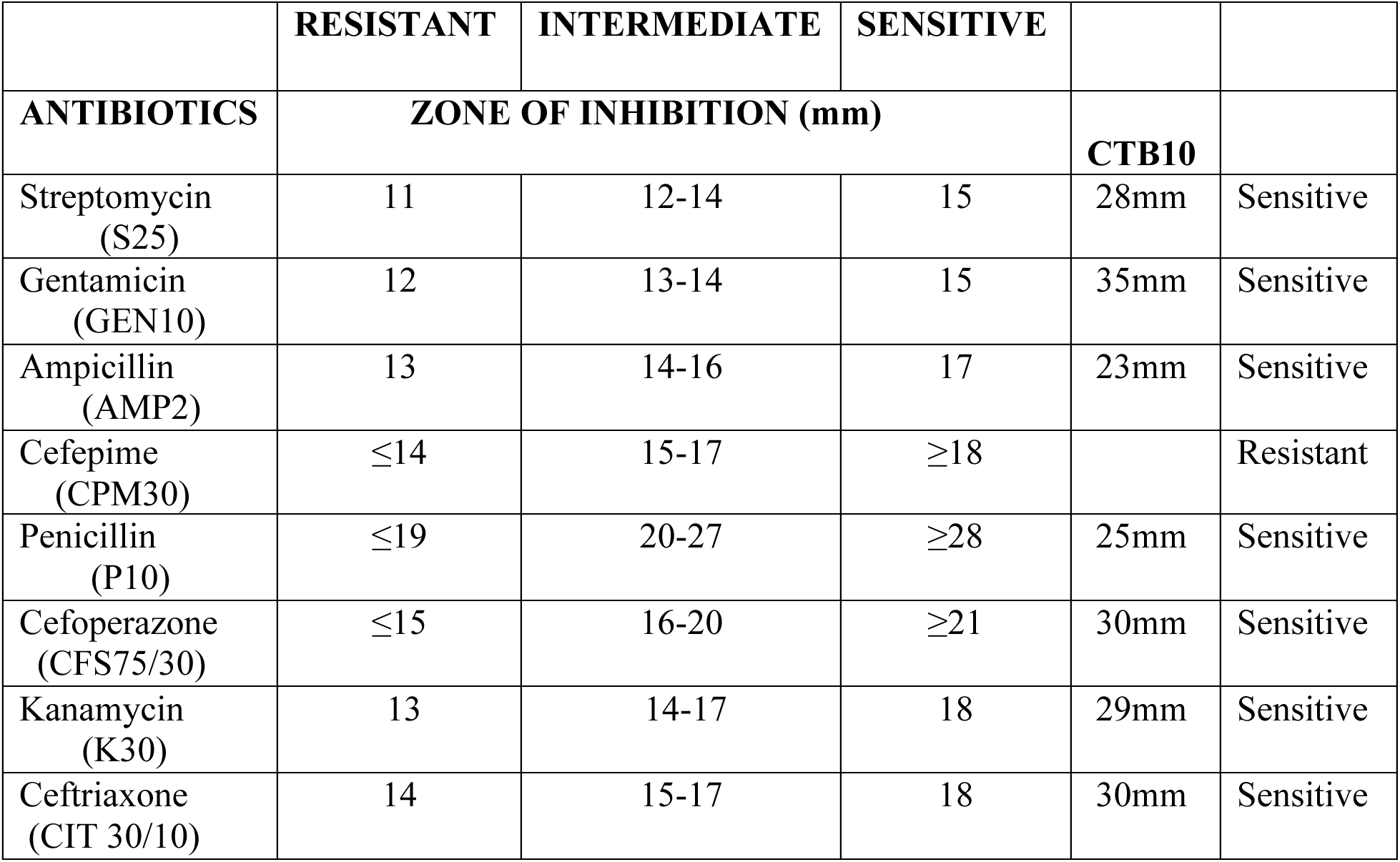
Antibiotic sensitivity test.

### MOLECULAR CHARACTERIZATION OF CTB10 STRAIN

16S rRNA sequence of CTB10 was searched in NCBI BLAST and it was confirmed that the CTB10 strain sequences shows matches with *Corynebacterium amycolatum* strain and the accession number PV258317 was obtained. Based on the earlier reports more than 50 distinct species are presently recognized as belonging to the genus *Corynebacterium*, which includes a remarkably wide variety of organisms. Still, it is widely known that the genus is not monophyletic and that it consists of multiple unique rDNA lineages (Chen et al.,2004). Many genera of *Corynebacterium amycolatum* strain was found in nature but each have different sequences and corelation. Mostly, it was a skin associated bacteria (Swaney et al.,2025) and rarely found in saline soil (Chen et al.,2004).

### PHYLOGENETIC ANALYSIS OF CTB10 STRAIN

The phylogenetic analysis of *Corynebacterium amycolatum* CTB10 with NR 024570.1 *Escherichia coli* strain U 5/41 acts as the outgroup. CTB10 has shown high similarity with *Corynebacterium amycolatum* strain CCUG35685 (Figure 4). The *Corynebacterium amycolatum* was not found to produce melanin earlier.

**Figure 4:**
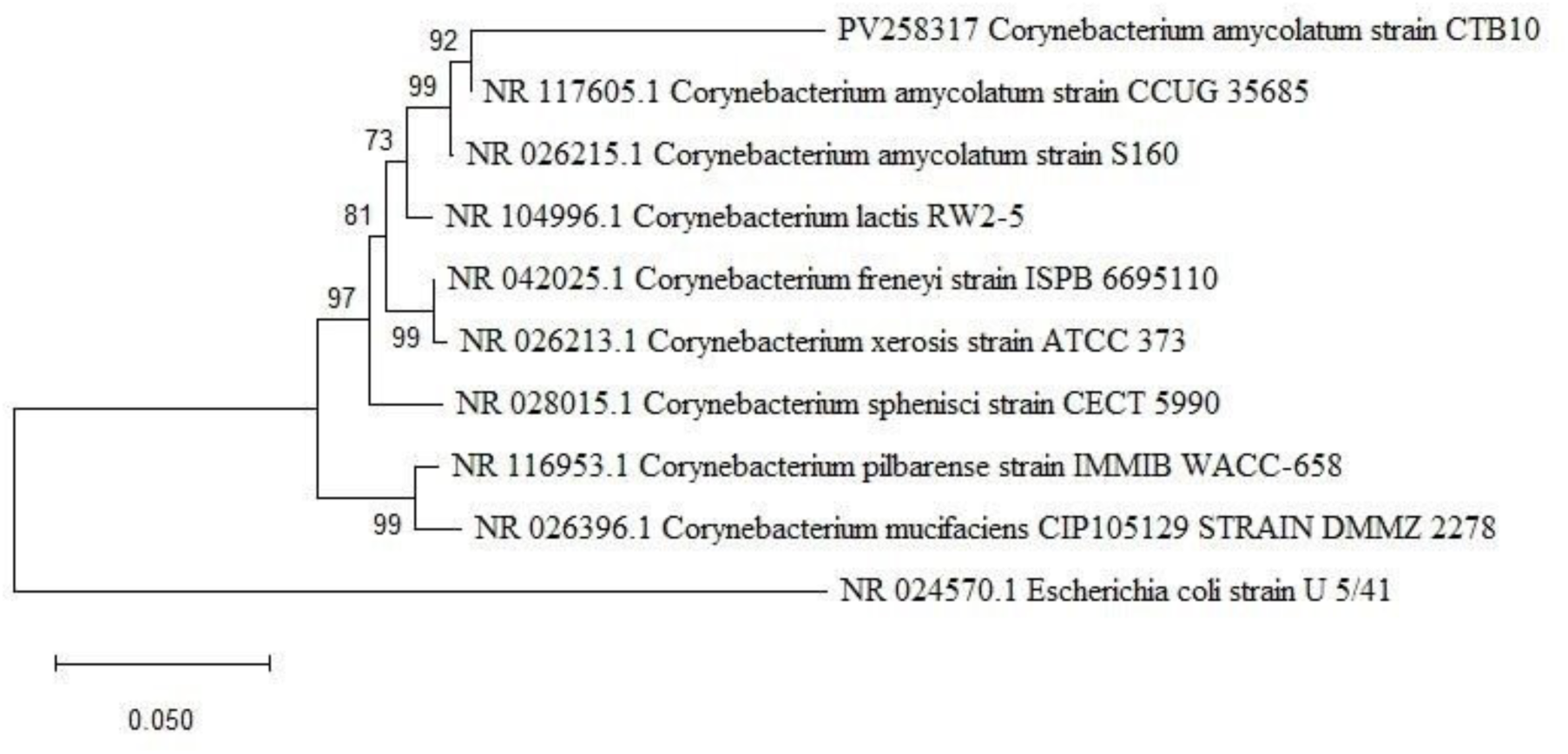
Phylogenetic tree of *Corynebacterium amycolatum* CTB10

### ANTIBIOTIC SENSITIVITY PROFILING

The CTB10 strain was showing sensitivity in most of the antibiotics tested. The CTB10 strain was only resistant to the Cefepime (CPM) disc (Figure 5b). Therefore, this result is like the earlier report of non-diphtheriae *Corynebacterium* isolates, and the commonly used antibiotic disc was ineffective of this strain (Neemuchwala et al., 2018). *Corynebacterium amycolatum* typically exhibits strong resistance to popular antibiotics such as gemifloxacin and ciprofloxacin, and telithromycin susceptibility is moderate (Hernández et al.,2003). The MAR index for CTB10 strain was found to be 0.125 which indicate the strain is isolated from less antibiotic polluted area and the strain CTB10 could be used for industrial production of melanin.

**Figure 5:**
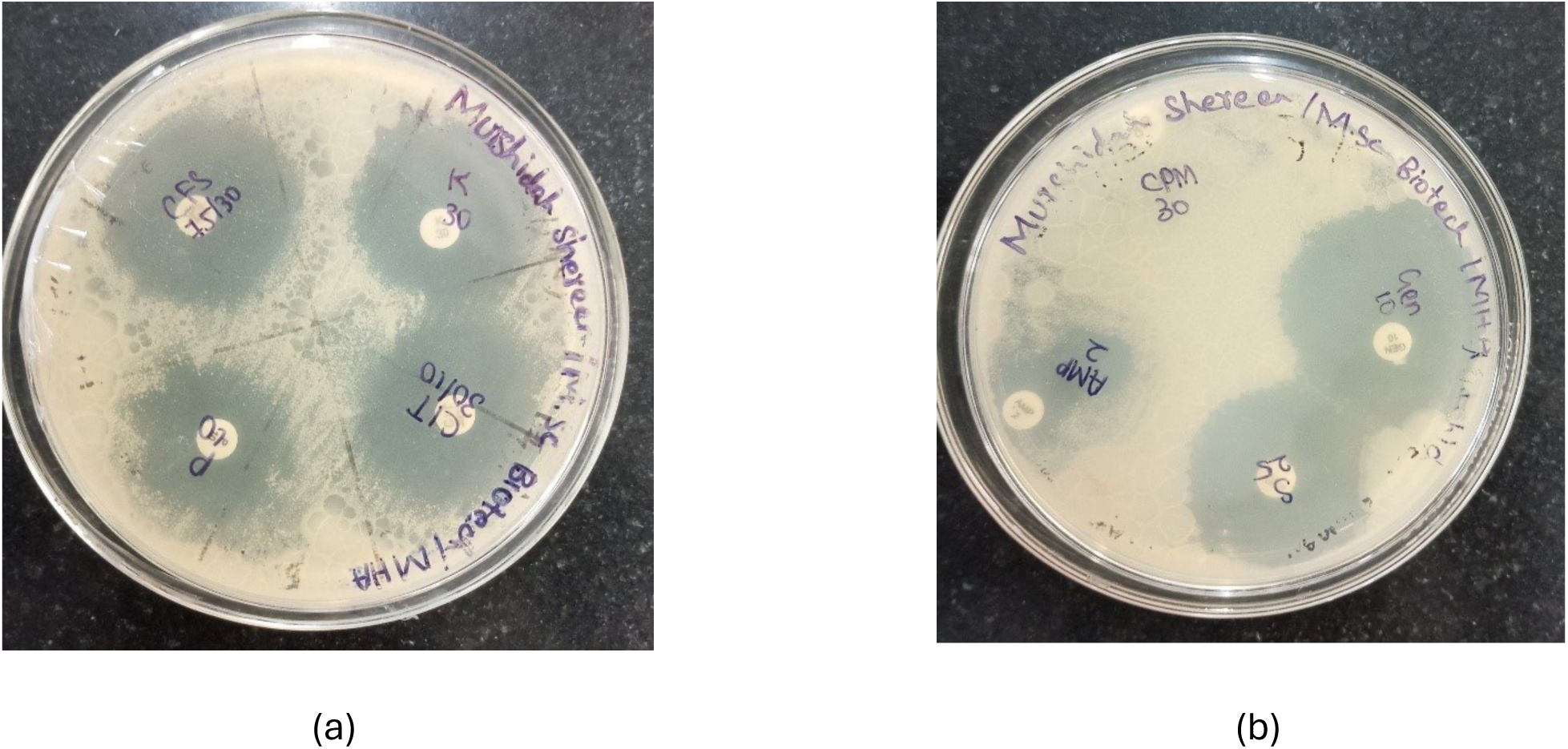
Cefoperazone, Kanamycin, Ceftriaxone and Penicillin disc and its zone of inhibition shown in (Figure 5a). Ampicillin, Streptomycin, Gentamicin disc has clear zone of inhibition and Cefepime disc alone resistant against CTB10 shown in (Figure 5b)

### HALOTOLERANCE OF STRAIN CTB10

The CTB10 strain was observed to exhibit halotolerance. The results of a 24-hour incubation period at 37°C indicate that the CTB10 strain exhibits some growth up to 10% NaCl, indicating that it possesses moderate halotolerant properties (Figure 6). According to previous research, high salt content stimulates the melanin production in most of the melanin producing bacteria. Melanin producing bacteria were found in the salt desert of Kutch, Gujarat (Jigna et al.,2022) and marine sediments (Kurian & Bhat,2018).

**Figure 6:**
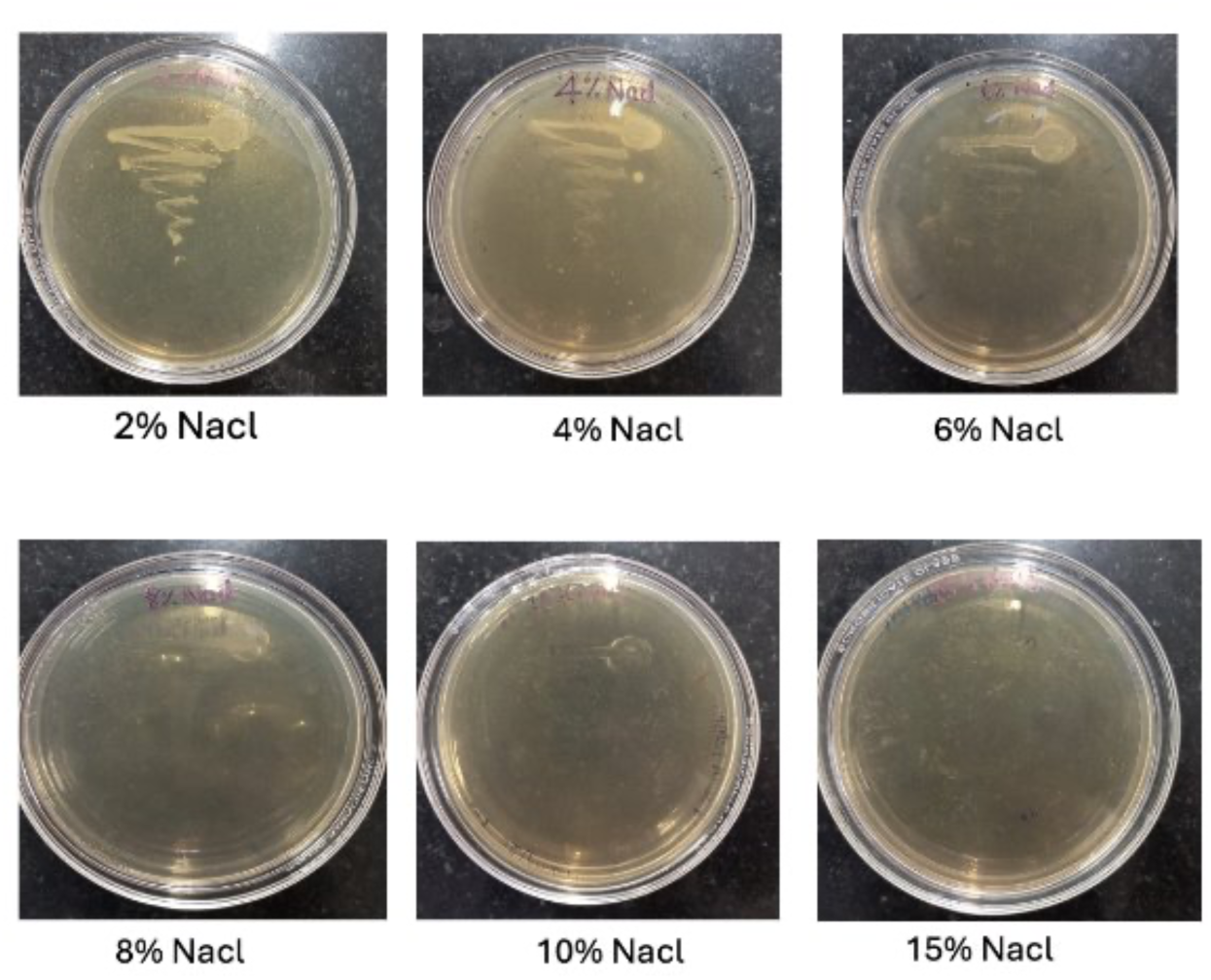
Halotolerance Assay

### EXTRACTION AND PURIFICATION OF MELANIN

After the TBB medium was turned into a dark colour, the melanin precipitated at pH 2, and the precipitation was clearly visible. The precipitated melanin was washed multiple times with solvents such as 0.1N HCL, 70% ethanol and distilled water. The purified powdered melanin used for further characterization. The most common methods for purifying melanin are acid hydrolysis and a series of washing procedures using organic solvents including acetone, petroleum ether, chloroform, ethyl acetate, or pure ethanol (Pralea et al.,2019). The same pattern of melanin extraction and purification method was used in the *Pseudomonas aeruginosa* strain was isolated from clinical samples (Gheni & Odaa., 2023).

### CHARACTERIZATION OF MELANIN SOLUBILITY TEST

The most common method for the identification and confirmation of melanin is its solubility. Melanin was found to be soluble in dimethyl sulfoxide (DMSO) and so, but it is largely insoluble in water and other inorganic and organic solvents like the earlier reported bacterial melanin samples (Singh et al., 2025). (Table 3)

**Table 3:** Solubility test.

| S.NO | SOLVENTS | RESULTS |
| --- | --- | --- |
| 1. | Water | Insoluble |
| 2. | Ethanol | Insoluble |
| 3. | Methanol | Insoluble |
| 4. | Isopropanol | Insoluble |
| 5. | Hydrochloric acid | Insoluble |
| 6. | Sulfuric acid | Sparingly soluble |
| 7. | DMSO | Soluble |
| 8. | Sodium hydroxide | Soluble |

**Table 4:** SPF potential of CTB10 melanin.

| Wavelength | EE($\lambda$) × I($\lambda$) | F1 | | F2 | |
| --- | --- | --- | --- | --- | --- |
| | | Abs | Abs × EE($\lambda$) × I($\lambda$) | Abs | Abs × EE($\lambda$) × I($\lambda$) |
| 290 | 0.015 | 0.541 | 0.008115 | 0.412 | 0.00618 |
| 295 | 0.0817 | 0.535 | 0.0437095 | 0.420 | 0.034314 |
| 300 | 0.2874 | 0.524 | 0.1505976 | 0.419 | 0.1204206 |
| 305 | 0.3278 | 0.509 | 0.1668502 | 0.399 | 0.1307922 |
| 310 | 0.1864 | 0.491 | 0.0915224 | 0.376 | 0.0700864 |
| 315 | 0.0837 | 0.473 | 0.0395901 | 0.360 | 0.030132 |
| 320 | 0.018 | 0.479 | 0.008622 | 0.328 | 0.005904 |
|  | **Total** |  | 0.5090068 |  | 0.3978292 |
|  | **SPF** |  | 5.090068 |  | 3.978292 |
| **CTB 10 strain had the SPF value 4.53 ± 0.7** |  |  |  |  |  |

### UV- VISIBLE SPECTROPHOTOMETRY

In UV-visible spectroscopy determination reveals that CTB10 has no characteristic absorbance peaks. The absorbance was high at 222nm and gradually decreases at 800nm in the UV-visible spectrum. Most of the melanin typically exhibits this kind of absorbance without characteristic peaks (Figure 7). This is because melanin lacks unique absorption peaks that set it apart from other cutaneous chromophores. As wavelengths rise from 300 to 800 nm, however, melanin monotonically declines in absorption (Ammanagi et al., 2021). This has happened due to the complex conjugated molecules in the melanin structure, which absorb and scatter UV light photons, are the cause of melanin’s high UV light absorption (Pralea et al.,2019).

**Figure 7:**
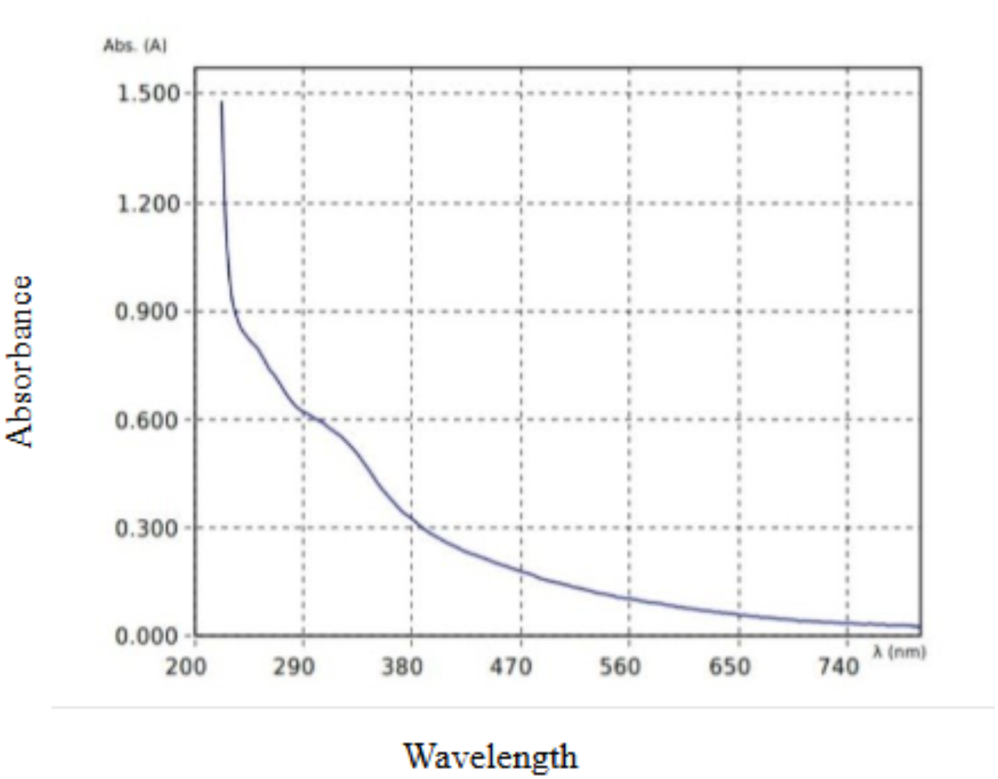
UV-Visible spectrum of CTB10 melanin showing the characteristics pattern

### FT-IR SPECTROPHOTOMETRY

CTB10 melanin shows the notable FT-IR spectrum are 3273.57cm^-1^, 1624.73cm^-1^, 1514.31cm^-1^ and 554.434cm^-1^ (Figure 8). The large absorbance 3273.57cm^-1^, which was due to O-H stretching or N-H stretching vibration of carboxylic acid and phenol group in the melanin. This FT-IR spectrum was like the earlier reports of Kiran et al., 2017 and the strong band at 1623 cm^-1^ is typical and was due to vibrations of the COO-groups or the aromatic ring C=C of the amide C=O. Aliphatic C-H groups may be responsible for bands at approximately 1400 to 1500 cm^-1^, while alkene C-H substitution in the melanin pigment is responsible for weak bands below 700 cm^-1^ (Tarangini & Mishra.,2013). Therefore, FT-IR analysis confirmed CTB10 strain produced pigment was melanin because it possesses the similar FT-IR peaks (Pralea et al.,2019).

**Figure 8:**
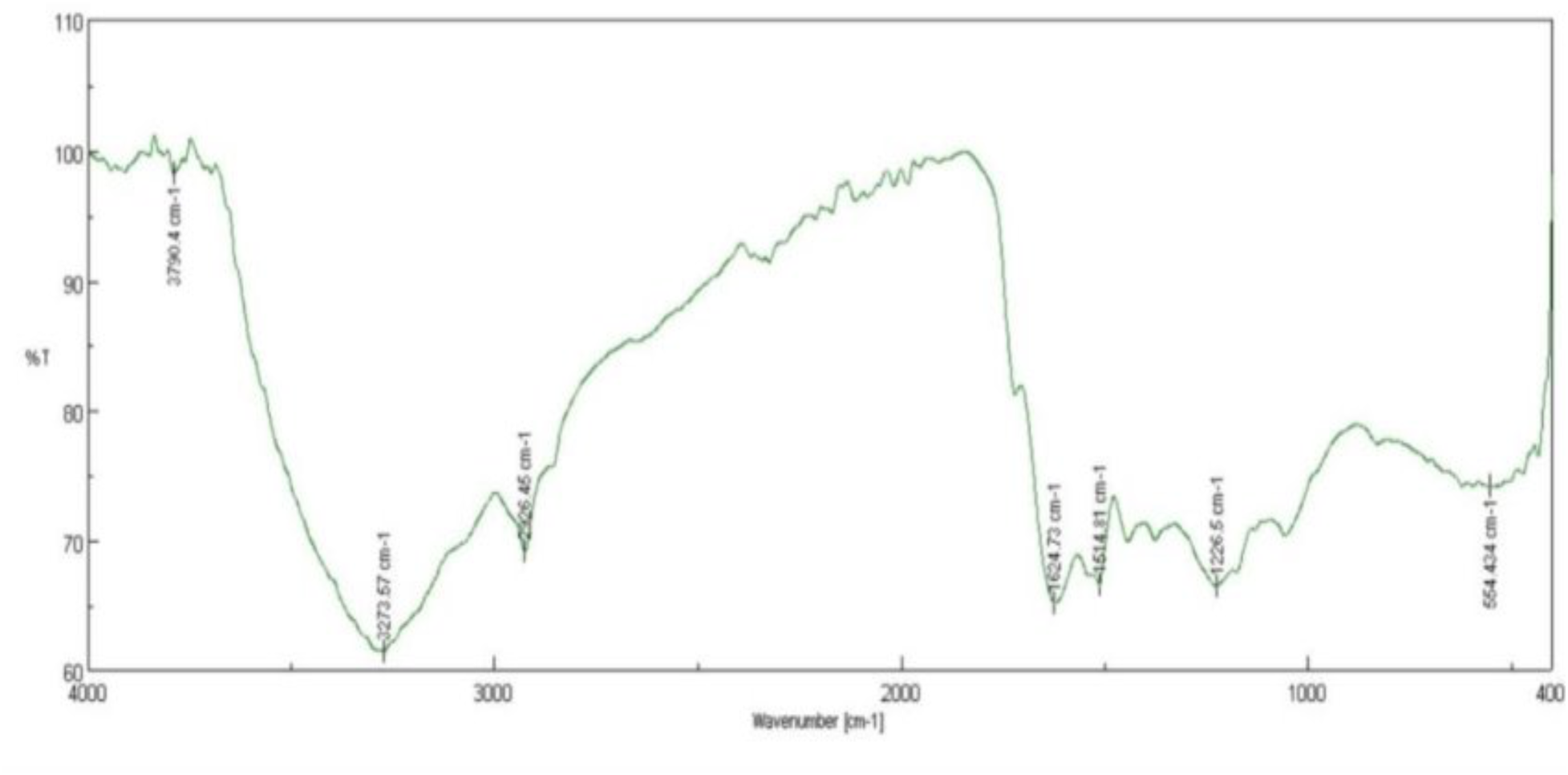
FT-IR spectrum of CTB10 melanin

### FIELD EMISSION SCANNING ELECTRON MICROSCOPY (FE-SEM)

The FE-SEM analysis was performed to investigate the morphological characteristics of the melanin nanoparticles. As shown in Figure 9A, the nanoparticles exhibited a highly aggregated structure at a magnification of 50,000×. This aggregation may be attributed to strong intermolecular interactions such as hydrogen bonding or π–π stacking, which are commonly observed in melanin-like materials due to their abundant aromatic and functional groups. The spherical agglomeration of the nanostructure observed in the surface morphology of melanin nanoparticles is consistent with earlier reports (Alcalá-Alcalá et al., 2023). The inset image (Figure 9B) provides a magnified view of a selected region, highlighting the nanoscale roughness and fine particulate texture of the material. As seen in the figure, the grains are spherical with ∼ sub-100 nm diameter and possibly are individual or loosely bound nanoparticles within the aggregate mass. These features suggest that while primarily the particles exist in the nano range of size, they tend to have clustered during or after synthesis. This indicates further need for surface passivation in order to improve colloidal stability and prevent particle agglomeration (Chen et al., 2025).

**Figure 9.**
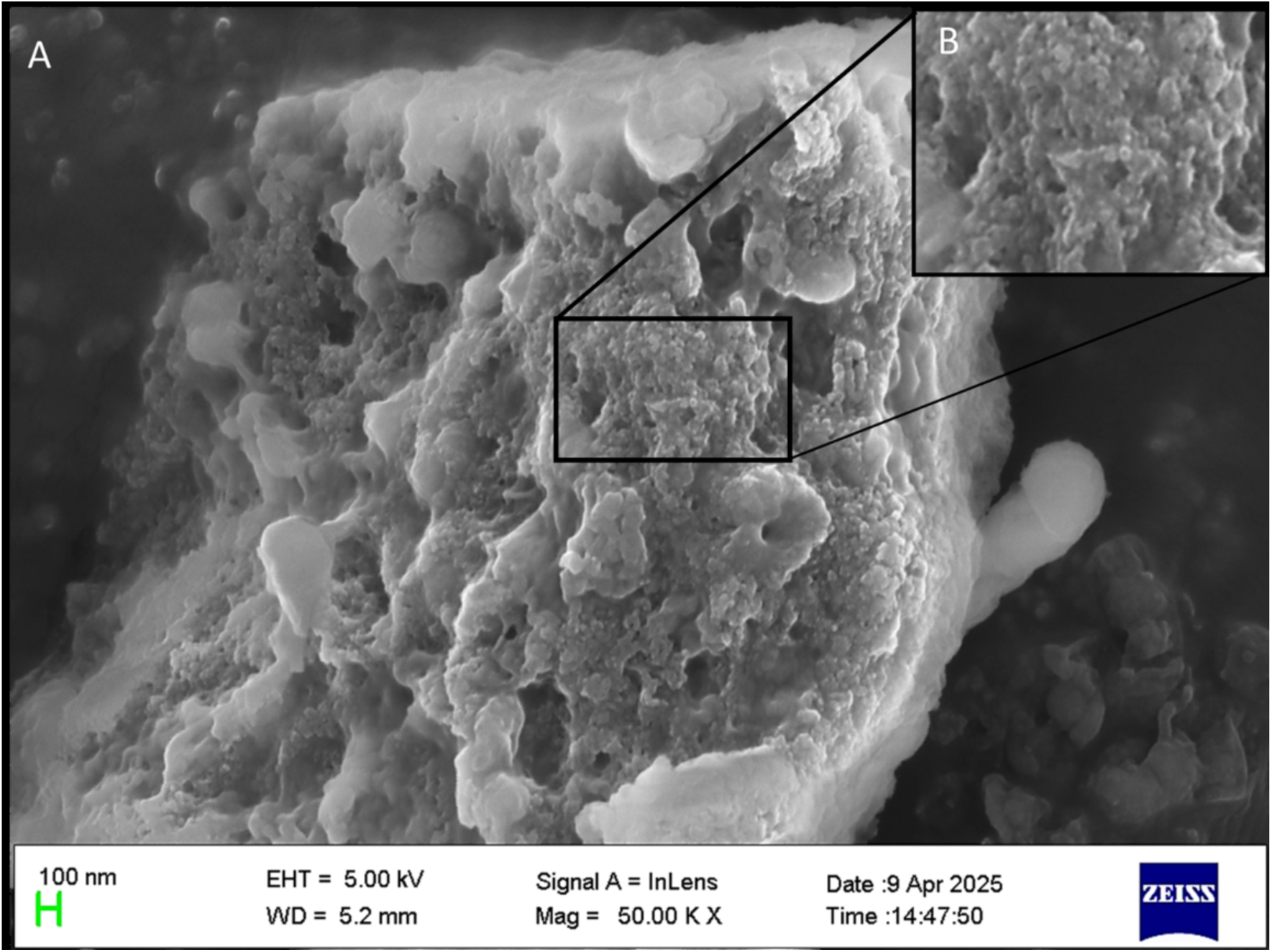
FE-SEM images of melanin nanoparticles. (a) Aggregated melanin nanoparticles under 50,000X magnification. (b) Callout shows a zoomed-in image of the selected region.

### COSMETIC PROPERTY OF THE MELANIN

#### SUN PROTECTION FACTOR (SPF) ESTIMATION

An *in vitro* spectrophotometric technique was utilized to determine SPF (Patil et al., 2016). The Sun Protection Factor (SPF), which is the ratio of the UV energy needed to produce a minimal erythemal dose (MED) in protected skin to the UV energy needed to produce a MED in unprotected skin, is typically used to represent how effective sunscreens are. The SPF values were calculated using the UV spectrophotometric method using the Mansur equation (Patki et al.,2021). The SPF value of CTB10 melanin by an average of 4.53 ±0.7. Based on the previous research SPF value of melanin *Amorphotheca resinae* SPF value 2.5, *Vibrio alginolyticus* strain BTKKS3 SPF value 3.42 (Kurian & Bhat,2018). The microbial melanin may be a good option for a bioinspired sunscreen due to its industrial scale and environmental sustainability. It is very biocompatible and has no negative effects on the environment after disposal. For many years, sunscreens have been used to protect people against UV rays, particularly UV-A (320–400 nm) and UV-B (280–320 nm). The bioactive components found in melanin are preferred as main elements in cosmetics because of their natural anti-mutagenic, anti-cancerous, and non-toxic qualities, since synthesized chemical photoprotective substances are more likely to be harmful and carcinogenic (http://www.jetir.org/papers/JETIRFY06006.pdf). The strain CTB10 melanin had a better SPF value compared to earlier reports therefore, the melanin could be used in cosmetic formulation after some optimization of SPF.

### ANTIOXIDANT ACTIVITY OF CTB10 MELANIN DPPH RADICAL SCAVENGING ASSAY

Antioxidants are used as active components in a lot of cosmetic products. The oxidation reactions that are triggered by environmental pollution might result in free radicals, which may be triggered by a series of events that harm skin cells. Finally, it causes skin-related problems, like dryness, pigmentation, wrinkles, and aging, are increasing free radicals. By eliminating the free radical intermediates and oxidizing themselves, which was done by cosmetic antioxidants able to stop chain reactions and prevent further oxidation processes, protecting the skin from the environmental stress that free radicals generate (Kusumawati, & Indrayanto.,2013). The antioxidant property ofvmelanin was immense when compared to the standard antioxidant ascorbic acid. The antioxidant property is widely used in most of the cosmetic products in the market, but bacterial melanin has this property naturally. This makes bacterial melanin was attractive replacement for chemically synthesized one. The naturally synthesized CTB10 melanin antioxidant activity was analysed by DPPH method. The result shows, there was a gradual increase in antioxidant efficacy when increasing the concentration of the melanin. The CTB10 melanin has the highest antioxidant value 61.25% of radical scavenging activity, with an IC50 value was 569.47µg/ml, and standard ascorbic acid highest antioxidant value 94.94% of radical scavenging activity with an IC50 28.19µg/ml (Figure 10). A higher amount of melanin has been associated with higher antioxidant activity, according to earlier research (Polapally et al., 2022).

**Figure 10:**
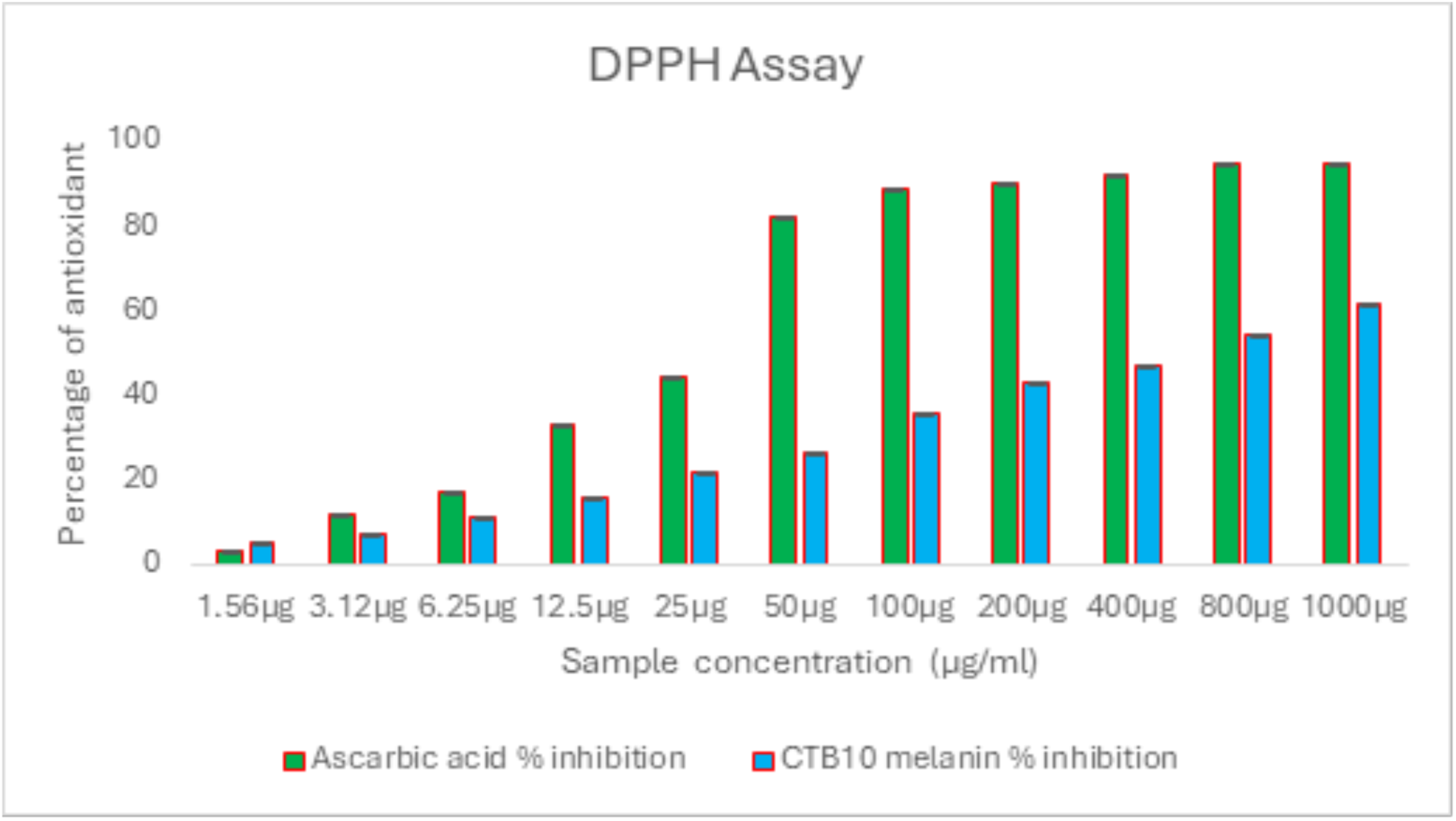
Radical scavenging activity of CTB10 melanin

### CYTOTOXICITY OF MELANIN

The cytotoxicity of CTB10 melanin shows very less cytotoxicity against the L929 mouse fibroblast cell line. Cell viability was determined to be 91.67% even when the melanin concentration was increased (Figure 11). According to earlier reports bacterial melanin was test in various cell lines and the outcome of the results proved melanin was non-cytotoxicity and some previous reports was BTCZ31 melanin was discovered to have a cytotoxic concentration of 105.4 μg/ml (IC50), which reduced the development of the L929 cell line (Kurian et al.,2015). Melanin pigment has the potential to be used as a natural anticancer agent due to its strong cytotoxic activity against the HFB4 skin cancer cell line and low cytotoxicity against healthy non-cancerous cells. *Streptomyces glaucescens* strain NEAE-H developed a melanin pigment that exhibited strong cytotoxic activity against the HFB4 skin cancer cell line (El-Naggar, N. E. A., & El-Ewasy, S. M. (2017). Melanin functions as a strong anticancer agent, killing cancer cells at low doses without harming healthy cells. Due to its significant anticancer activity against the SK-MEL-28 cell line and low cytotoxicity to healthy, noncancerous cells. The melanin pigment may be employed as a possible antitumor drug (Polapally et al.,2022). Therefore, these were confirmed bacterial melanin was safe to be used in appropriate dose.

**Figure 11:**
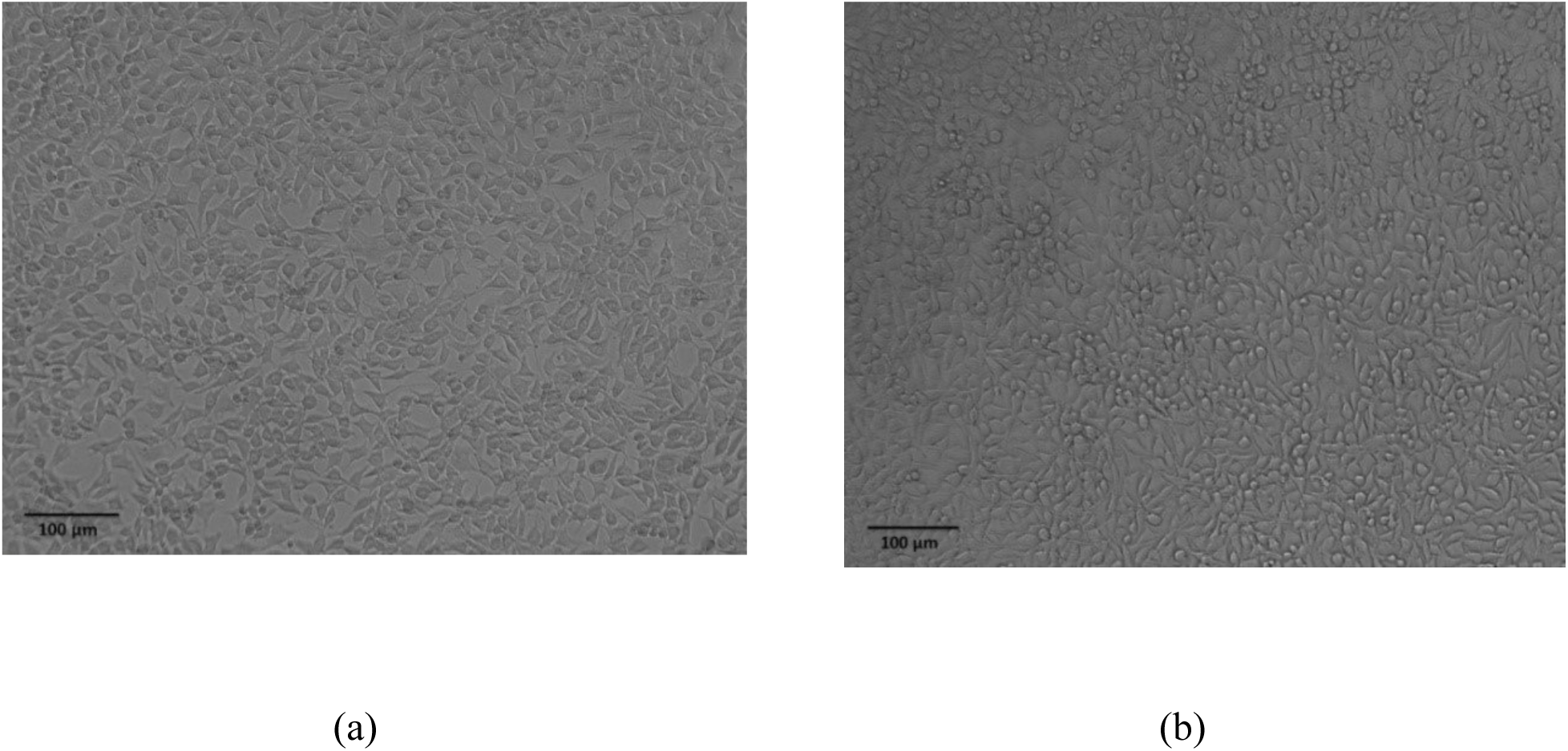
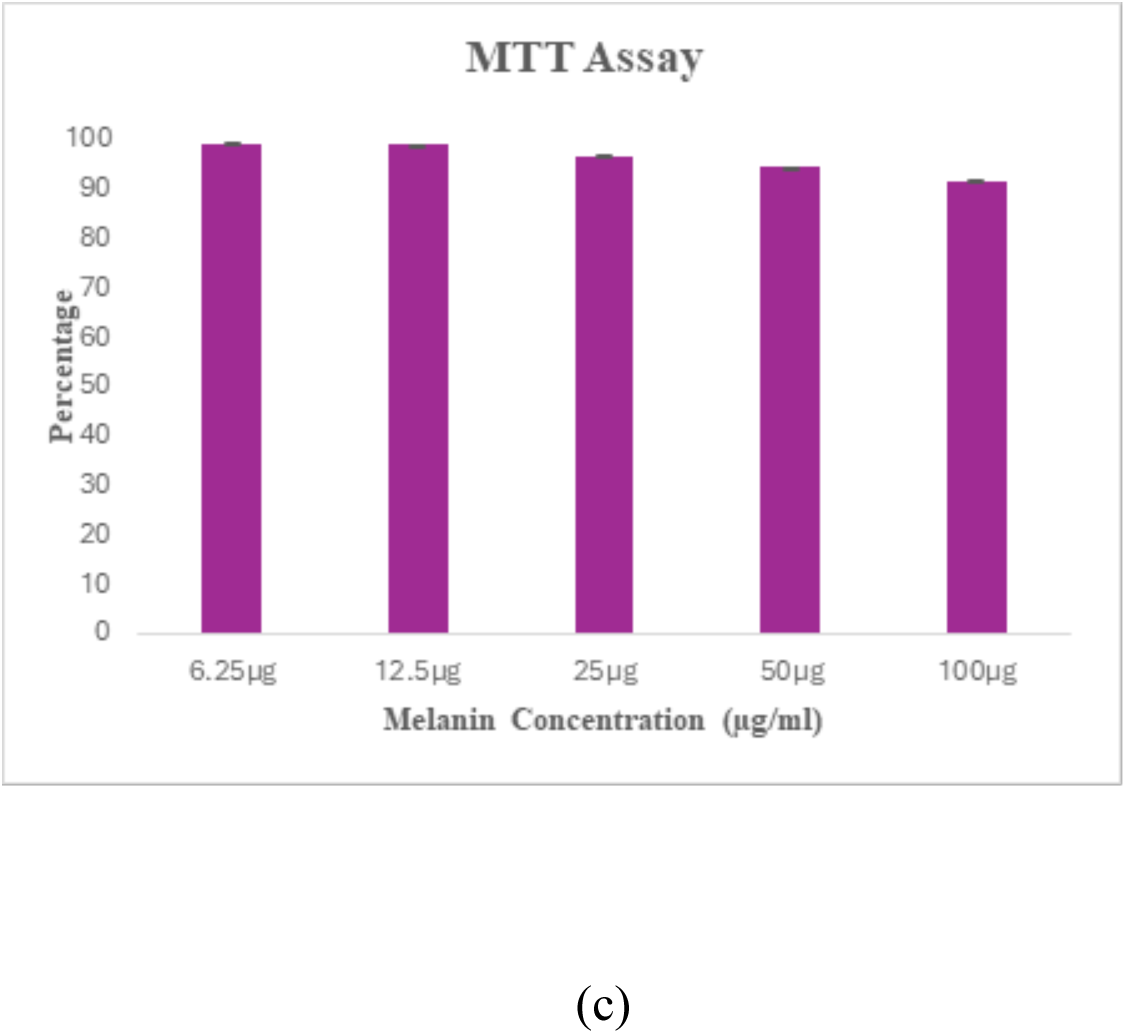
MTT assay (a) control (b) 100µg melanin higher concentration (c) Percentage viability graph

### *IN-VIVO* TOXICITY STUDY OF ZEBRA FISH EMBRYO

The toxicity study of zebra fish embryos tested by CTB10 melanin shows that the phenotypic variation of all embryos same to that of the control embryos (untreated melanin well) without any cause for the embryo. The 100% percentage of zebra fish embryo viability was noted in 24 hours of incubation, and a slight death rate was noted on 48 hours (Figure 12b). The CTB10 melanin does not affect the hatching of embryos and there no malformation of embryo was noted. This analysis confirms CTB10 melanin does not affect the delay of hatching and phenotype of zebra fish embryo (Figure 12a). Most of the earlier reports use zebra fish embryo studies to detect the toxicity of the plant derived compounds. There are two techniques to classify and assess developmental toxicity in zebrafish embryos in comparison with control embryos. First, coagulation, heartbeat absence, and tail separation are used to determine death. Second, aberrant eye development, absence of pigmentation, and lack of spontaneous movement are used to assess fetal development. Normal development has been proven by the control (Chahardehi et al.,2020).

**Figure 12:**
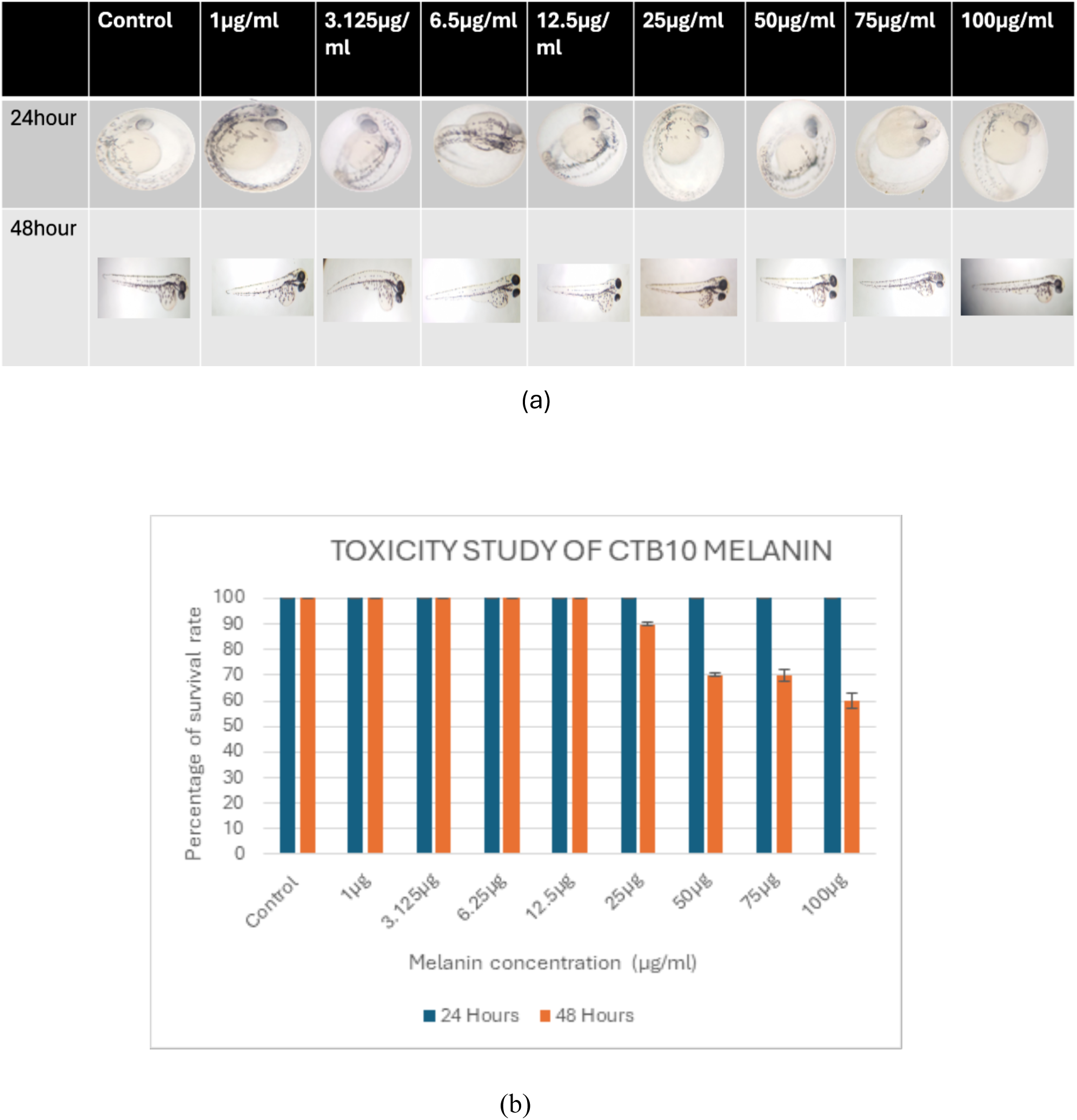
*in-vivo* toxicity study – (a) Microscopic images of zebra fish embryo in different time periods (b) toxicity effect on CTB10 melanin

### STUDYING MELANIN METABOLISM IN STRAIN CTB10 USING INHIBITORS

CTB10 melanin production was significantly inhibited by kojic acid in a concentration depended on manner. In visual observation could be clearly shown 2mM kojic acid concentration TBB tube of brown colour faded away than the control TBB tube (Figure 13a). This means when increase the concentration of kojic acid fades the original colour of bacterial melanin. The spectrophotometrically the concentration melanin produced in kojic acid treated tubes were compared to the untreated control. It was found control has produced 535.02µg/ml melanin which kojic acid at higher concentration produced 247.9µg/ml of melanin. So, the kojic acid inhibits the CTB10 melanin biosynthesis pathway and it proved CTB10 strain predominantly used the DOPA pathway for melanin synthesis. In an earlier report of inhibitor analysis of melanin was tested in *Klebsiella sp.*GSK and it also confirms the strain melanin was significantly inhibited by kojic acid (Sajjan et al., 2010) as like CTB10 strain. Some previous reports explain the working of kojic acid in melanin inhibition and the reports were Potential suppression of cellular NF-κB activity in human keratinocytes was demonstrated by kojic acid. The copper ion was captured by Kojic acid, which stops it from activating the tyrosinase; this was the working mechanism, and it leads to stopping the melanin production. This kojic acid is widely used in skin care cosmetic products to reduce melanin production (Saeedi et al.,2019) and two well-known tyrosinase inhibitors and depigmenting agents are kojic acid and β-arbutin. These are frequently used as positive controls to screen for new substances or extracts that successfully prevent the formation of melanin. This previous research paper tested the inhibitor activity of kojic acid, α-arbutin, β-arbutin, and deoxyarbutin on tyrosinase from mushrooms and in B16F10 mouse melanoma cells, and the outcome of results show that kojic acid consistently blocks melanin than β-arbutin. So, kojic acid was a better skin lightening agent (Wang et al.,2022).

**Figure 13:**
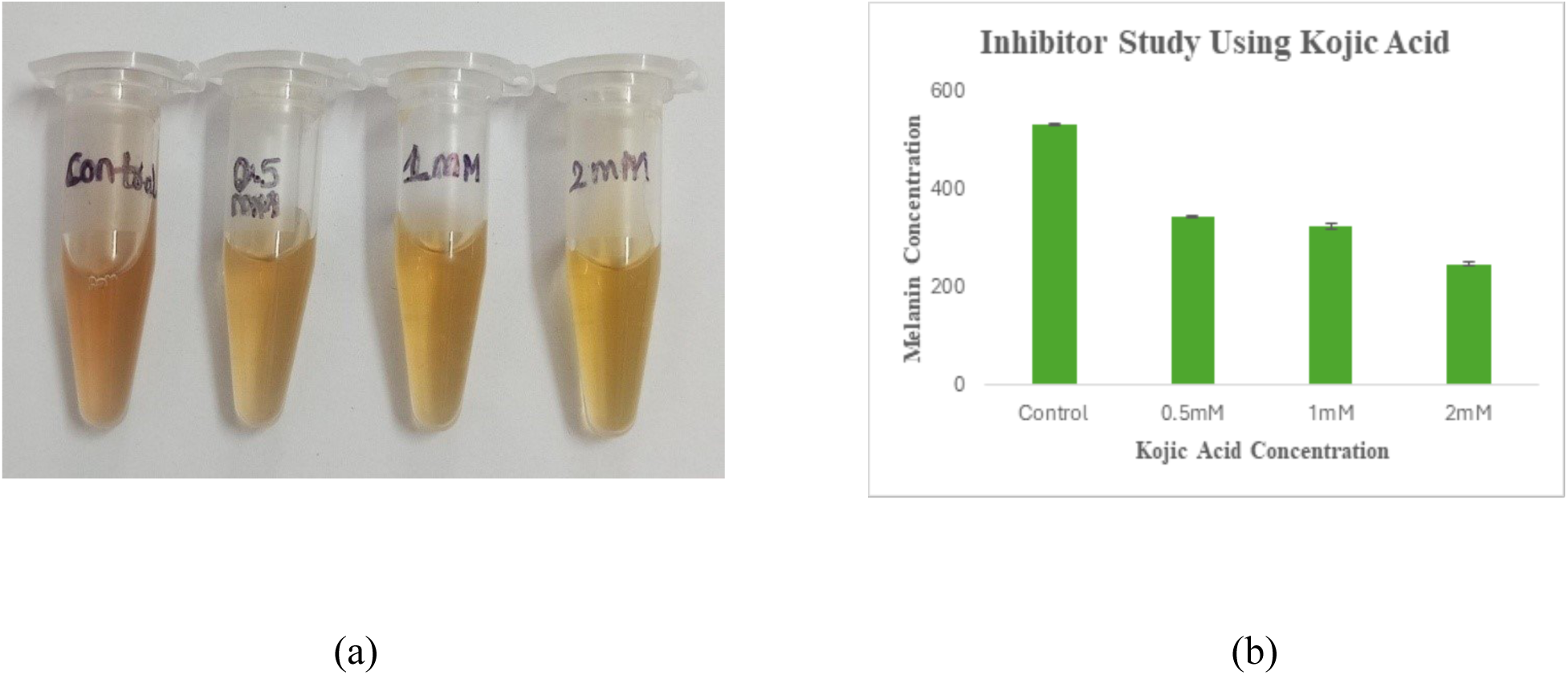
Inhibitor study using kojic acid (a) colour fade CTB10 melanin after centrifuge (b) melanin concentration graph (melanin concentration: control- 535.02µg/ml, 0.5mM-346.12µg/ml, 1mM-326.04µg/ml and 2mM- 247.9µg/ml).

## CONCLUSION

According to the results shown above, the novel report of *Corynebacterium amycolatum* CTB10 was a fast melanin producing bacterium. Characteristics of melanin was confirmed, as the produced pigment exactly matched with already existing bacterial melanin properties based on an earlier reports. The *Corynebacterium amycolatum* CTB10 melanin has cosmetic applications, which were confirmed by its sun protection factor, good antioxidant activity and less cytotoxicity. The inhibitor study confirmed that *Corynebacterium amycolatum* CTB10 melanin falls under the DOPA pathway for melanin biosynthesis. Cytotoxicity observation is necessary to prove that the melanin is a safe ingredient in the cosmetic product.

## Notes

### Competing Interest Statement

The authors have declared no competing interest.

## REFERENCES

Nosanchuk, J. D., & Casadevall, A. (2003). The contribution of melanin to microbial pathogenesis. Cellular microbiology, 5(4), 203–223.

Pavan, M. E., López, N. I., & Pettinari, M. J. (2020). Melanin biosynthesis in bacteria, regulation and production perspectives. Applied Microbiology and Biotechnology, 104(4), 1357–1370.

Singh, R., Rahman, S., Abbas, S.A. et al. Melanin: Comprehensive Insights into Pathways and Sustainable Applications. Curr. Pharmacol. Rep. 11, 37 (2025).

El-Zawawy, N. A., Kenawy, E. R., Ahmed, S., & El-Sapagh, S. (2024). Bioproduction and optimization of newly characterized melanin pigment from Streptomyces djakartensis NSS-3 with its anticancer, antimicrobial, and radioprotective properties. Microbial Cell Factories, 23(1), 23.

Jigna, C., Mira Gordhanbhai, P., & Kurian K. N. (2022). Bacterial Melanin with Immense Cosmetic Potential Produced by Marine Bacteria Bacillus pumilus MIN3. Preprints. 10.20944/preprints202205.0296.v1

Choi, K. Y. (2021). Bioprocess of microbial melanin production and isolation. Frontiers in bioengineering and biotechnology, 9, 765110.

Hemraj, V., Diksha, S., & Avneet, G. (2013). A review on commonly used biochemical test for bacteria. Innovare Journal of Life Science, 1(1), 1–7.

Janda, J. M., & Abbott, S. L. (2007). 16S rRNA gene sequencing for bacterial identification in the diagnostic laboratory: pluses, perils, and pitfalls. Journal of clinical microbiology, 45(9), 2761–2764.

Calvopiña, M., Toro, M., Bastidas-Caldes, C., Vasco-Julio, D., & Muñoz, G. (2023). A fatal case of disseminated histoplasmosis by Histoplasma capsulatum var. capsulatum misdiagnosed as visceral leishmaniasis—molecular diagnosis and identification. Pathogens, 12(9), 1112.

Hudzicki, J. (2009). Kirby-Bauer disk diffusion susceptibility test protocol. American society for microbiology, 15(1), 1–23.

Mir, R., Salari, S., Najimi, M., & Rashki, A. (2022). Determination of frequency, multiple antibiotic resistance index and resistotype of Salmonella spp. in chicken meat collected from southeast of Iran. Veterinary medicine and science, 8(1), 229–236.

Ilyas, N., Mazhar, R., Yasmin, H., Khan, W., Iqbal, S., Enshasy, H. E., & Dailin, D. J. (2020). Rhizobacteria isolated from saline soil induce systemic tolerance in wheat (Triticum aestivum L.) against salinity stress. Agronomy, 10(7), 989.

Rahman, S. S., Siddique, R., & Tabassum, N. (2017). Isolation and identification of halotolerant soil bacteria from coastal Patenga area. BMC research notes, 10, 1–6.

Ferraz, A. R., Pacheco, R., Vaz, P. D., Pintado, C. S., Ascensão, L., & Serralheiro, M. L. (2021). Melanin: Production from cheese bacteria, chemical characterization, and biological activities. International journal of environmental research and public health, 18(20), 10562.

Pralea, I. E., Moldovan, R. C., Petrache, A. M., Ilieș, M., Hegheș, S. C., Ielciu, I., … & Iuga, C. A. (2019). From extraction to advanced analytical methods: The challenges of melanin analysis. International journal of molecular sciences, 20(16), 3943.

Rudrappa, M., Kumar, S., Kumar, R. S., Almansour, A. I., Perumal, K., & Nayaka, S. (2022). Bioproduction, purification and physicochemical characterization of melanin from Streptomyces sp. strain MR28. Microbiological Research, 263, 127130.

Tarangini, K., & Mishra, S. (2013). Production, characterization and analysis of melanin from isolated marine Pseudomonas sp. using vegetable waste. Res J Eng Sci, 2278, 9472.

Patki, J. M., Singh, S., Singh, S., Padmadas, N., & Dasgupta, D. (2021). Analysis of the applicative potential of pigments extracted from bacterial isolates of mangrove soil as topical UV protectants. Brazilian Journal of Pharmaceutical Sciences, 57, e19127.

Antony, T. M. P., Krishna, A. R., Jayalekshmi, S. K., Chockalingam, J., & Ramasamy, S. (2023). Isolation and Elucidation of Bacterial Melanin’s Sun Protection Factor (SPF) for Photoprotection in Cosmetics. J. Pure Appl. Microbiol, 17, 449–455.

Brand-Williams, W., Cuvelier, M. E., & Berset, C. L. W. T. (1995). Use of a free radical method to evaluate antioxidant activity. LWT-Food science and Technology, 28(1), 25–30

Anthony, K. P., & Saleh, M. A. (2013). Free radical scavenging and antioxidant activities of silymarin components. Antioxidants, 2(4), 398–407.

Mosmann, T. (1983). Rapid colorimetric assay for cellular growth and survival: application to proliferation and cytotoxicity assays. Journal of immunological methods, 65(1-2), 55–63.

Joseph, M. M., Aravind, S. R., Varghese, S., Mini, S., & Sreelekha, T. T. (2012). Evaluation of antioxidant, antitumor and immunomodulatory properties of polysaccharide isolated from fruit rind of Punica granatum. Molecular medicine reports, 5(2), 489–496

Joshi, V., & Katti, P. (2018). Developmental toxicity assay for food additive tartrazine using zebrafish (Danio rerio) embryo cultures. International Journal of Toxicology, 37(1), 38–44.

Manivasagan, P., Venkatesan, J., Senthilkumar, K., Sivakumar, K., & Kim, S. K. (2013). Isolation and characterization of biologically active melanin from Actinoalloteichus sp. MA-32. International journal of biological macromolecules, 58, 263-274.

Choi, Y. J., Rho, H. S., Baek, H. S., Hong, Y. D., Joo, Y. H., Shin, S. S., & Kim, J. M. (2014). Synthesis and biological evaluation of kojic acid derivatives as tyrosinase inhibitors. Bulletin of the Korean Chemical Society, 35(12), 3647–3650.

Chen, H. H., Li, W. J., Tang, S. K., Kroppenstedt, R. M., Stackebrandt, E., Xu, L. H., & Jiang, C. L. (2004). Corynebacterium halotolerans sp. nov., isolated from saline soil in the west of China. International journal of systematic and evolutionary microbiology, 54(3), 779–782.

Swaney, M. H., Henriquez, N., Campbell, T., Handelsman, J., & Kalan, L. R. (2025). Skin-associated Corynebacterium amycolatum shares cobamides. mSphere, 10(1), e00606–24.

Neemuchwala, A., Soares, D., Ravirajan, V., Marchand-Austin, A., Kus, J. V., & Patel, S. N. (2018). In vitro antibiotic susceptibility pattern of non-diphtheriae Corynebacterium isolates in Ontario, Canada, from 2011 to 2016. Antimicrobial agents and chemotherapy, 62(4), 10-1128.

Hernández, J. S., Peris, B. M., Guirao, G. Y., Zufiaurre, N. G., Bellido, J. M., Hernández, M. S., & Rodríguez, J. G. (2003). In vitro activity of newer antibiotics against Corynebacterium jeikeium, Corynebacterium amycolatum and Corynebacterium urealyticum. International journal of antimicrobial agents, 22(5), 492–496.

Kurian, N. K., & Bhat, S. G. (2018). Data on the characterization of non-cytotoxic pyomelanin produced by marine Pseudomonas stutzeri BTCZ10 with cosmetological importance. Data in brief, 18, 1889.

Gheni, M. R., & Odaa, N. H. (2023). Production, Extraction, and Purification of An Extracellular Melanin Pigment from Clinically Isolated Pseudomonas aeruginosa. Egyptian Journal of Hospital Medicine, 92(1).

Ammanagi, A., Ct, S., Badiger, A., & Ramaraj, V. (2021, September). Functional and structural characterization of melanin from Brevibacillus invocatus strain IBA. In Doklady Biological Sciences (Vol. 500, pp. 159-169). Pleiades Publishing

Kiran, G. S., Jackson, S. A., Priyadharsini, S., Dobson, A. D., & Selvin, J. (2017). Synthesis of Nm-PHB (nanomelanin-polyhydroxy butyrate) nanocomposite film and its protective effect against biofilm-forming multi drug resistant Staphylococcus aureus. Scientific reports, 7(1), 9167.

Alcalá-Alcalá, S., Casarrubias-Anacleto, J. E., Mondragón-Guillén, M., Tavira-Montalvan, C. A., Bonilla-Hernández, M., Gómez-Galicia, D. L., Gosset, G., & Meneses-Acosta, A. (2023). Melanin Nanoparticles Obtained from Preformed Recombinant Melanin by Bottom-Up and Top-Down Approaches. Polymers, 15(10), 2381. 10.3390/polym15102381

Chen, J., Yu, J., & Lin, S. (2025). Insight into dispersibility regulation on arsenic removal using magnetic iron oxide nanoparticles. Applied Surface Science, 695, 162939.

Polapally, R., Mansani, M., Rajkumar, K., Burgula, S., Hameeda, B., Alhazmi, A., … & Sayyed, R. Z. (2022). Melanin pigment of Streptomyces puniceus RHPR9 exhibits antibacterial, antioxidant and anticancer activities. PLoS One, 17(4), e0266676.

Kusumawati, I., & Indrayanto, G. (2013). Natural antioxidants in cosmetics. Studies in natural products chemistry, 40, 485–505.

Kurian, N. K., Nair, H. P., & Bhat, S. G. (2015). Evaluation of anti-inflammatory property of melanin from marine Bacillus spp. BTCZ31. Evaluation, 8(3).

El-Naggar, N. E. A., & El-Ewasy, S. M. (2017). Bioproduction, characterization, anticancer and antioxidant activities of extracellular melanin pigment produced by newly isolated microbial cell factories Streptomyces glaucescens NEAE-H. Scientific reports, 7(1), 1–19.

Modarresi Chahardehi, A., Arsad, H., & Lim, V. (2020). Zebrafish as a successful animal model for screening toxicity of medicinal plants. Plants, 9(10), 1345.

Wang, W., Gao, Y., Wang, W., Zhang, J., Yin, J., Le, T., … & Jiang, H. (2022). Kojic acid showed consistent inhibitory activity on tyrosinase from mushroom and in cultured B16F10 cells compared with arbutins. Antioxidants, 11(3), 502.

Saeedi, M., Eslamifar, M., & Khezri, K. (2019). Kojic acid applications in cosmetic and pharmaceutical preparations. Biomedicine & Pharmacotherapy, 110, 582–593.

